# Hepatobiliary manganese homeostasis is dynamic in the setting of illness in mice

**DOI:** 10.1101/2023.03.22.533688

**Authors:** Laxmi Sunuwar, Vartika Tomar, Asia Wildeman, Valeria Culotta, Joanna Melia

**Affiliations:** Department of Medicine, Division of Gastroenterology and Hepatology, Johns Hopkins University School of Medicine, Baltimore, MD 21205, USA; Department of Biochemistry and Molecular Biology, Johns Hopkins University Bloomberg School of Public Health, Baltimore, MD 21205, USA

**Author notes:** **Correspondence:** Joanna Melia, MD, Johns Hopkins University School of Medicine, 720 Rutland Avenue, Ross 912, Baltimore, MD 21205. contributed equally to this work (Laxmi Sunuwar, Vartika Tomar).

**Keywords:** Manganese, colitis, inflammation, immune, liver, bile, mice, Slc39a8, Zip8

## Abstract

Manganese is a diet−derived micronutrient that is essential for critical cellular processes like redox homeostasis, protein glycosylation, and lipid and carbohydrate metabolism. Control of Mn availability, especially at the local site of infection, is a key component of the innate immune response. Less has been elucidated about Mn homeostasis at the systemic level. In this work, we demonstrate that systemic Mn homeostasis is dynamic in response to illness in mice. This phenomenon is evidenced in male and female mice, mice of two genetic backgrounds (C57/BL6 and BALB/c), in multiple models of acute (dextran−sodium sulfate−induced) and chronic (*enterotoxigenic Bacteriodes fragilis*) colitis, and systemic infection with *Candida albicans*. When mice were fed a standard corn−based chow with excess Mn (100 ppm), liver Mn decreased and biliary Mn increased 3−fold in response to infection or colitis. Liver iron, copper, and zinc were unchanged. When dietary Mn was restricted to minimally adequate amounts (10ppm), baseline hepatic Mn levels decreased by approximately 60% in the liver, and upon induction of colitis, liver Mn did not decrease further, however biliary Mn still increased 20−fold. In response to acute colitis, hepatic *Slc39a8* mRNA (gene encoding the Mn importer, Zip8) and *Slc30a10* mRNA (gene encoding the Mn exporter, Znt10) are decreased. Zip8 protein is decreased. Illness− associated dynamic Mn homeostasis may represent a novel host immune/inflammatory response that reorganizes systemic Mn availability through differential expression of key Mn transporters with down−regulation of Zip8.

**Graphical Abstract:** 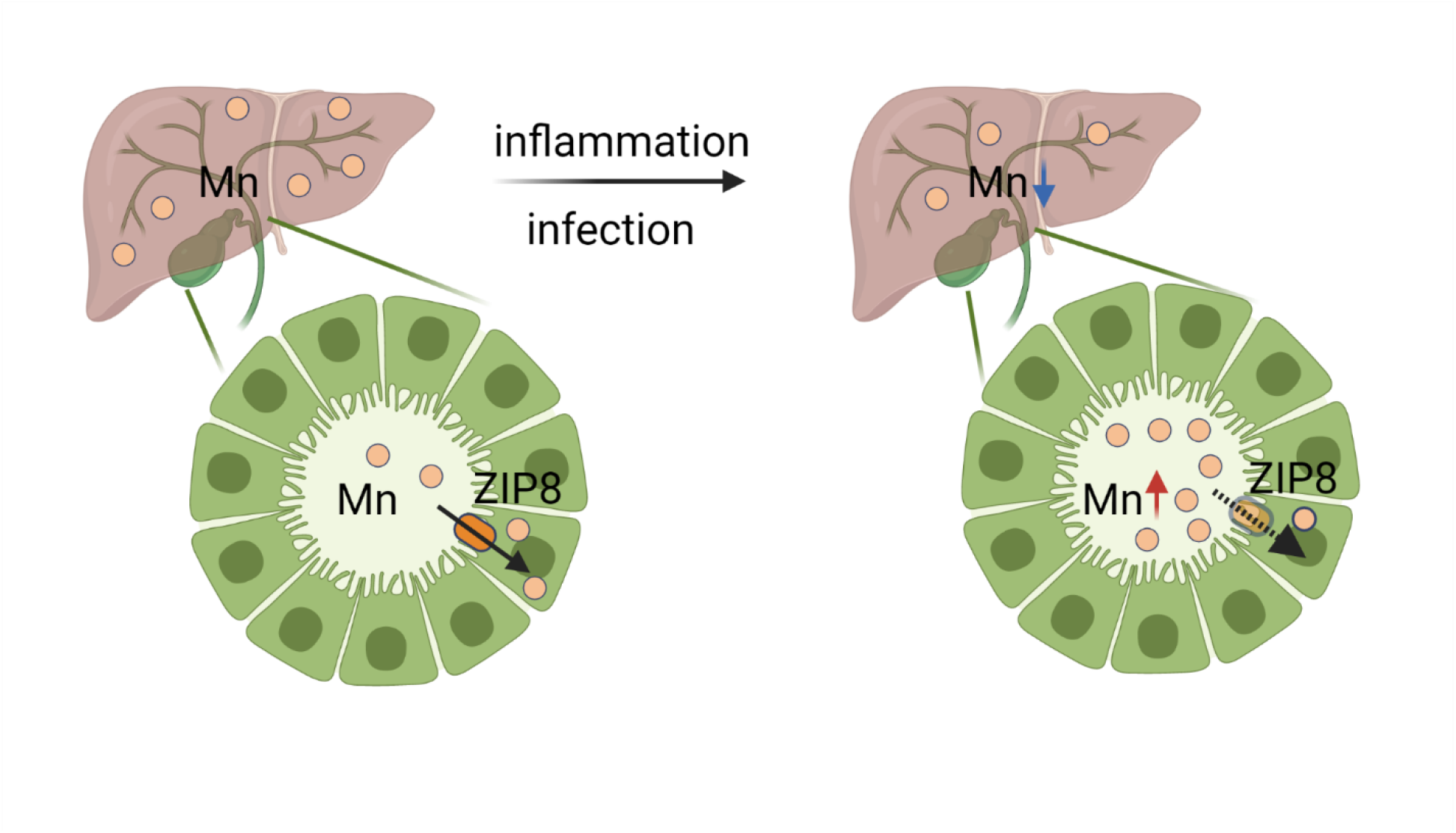

Created with BioRender.com

## Introduction

Transition metals including iron, zinc, copper, and manganese are important to support physiologic cellular function, with more than 30% of proteins interacting with metal cofactors. Of these transition metals, Mn is least abundant, but Mn is a critical cofactor for multiple metalloenzymes needed for cellular homeostasis, including redox homeostasis through the activity of Mn superoxide dismutase, lipid and carbohydrate metabolism, and protein glycosylation via metal requiring glycosyltransferases (1, 2). Through genome wide association studies and exome sequencing studies, there is an increasing awareness that aberrant Mn homeostasis contributes to risk for many chronic diseases (3), however many questions remain about how Mn is sensed, how the activities of the primary Mn transporters are coordinated, and, if and how Mn homeostasis is dynamic. In this work, we address this last question and present evidence that Mn homeostasis is dynamic in response to ‘illness’.

Human and mouse studies have elucidated that the liver is a key site of regulation of systemic Mn homeostasis. There are three main Mn transporters (ZIP14, ZNT10, and ZIP8) (4). Conceptually, ZIP14 (encoded by the gene *SLC39A14*) imports Mn into the liver from blood, while ZNT10 (encoded by *SLC30A10*) actively exports Mn across the bile canalicular membrane for excretion into bile and ZIP8 (encoded by *SLC39A8*) serves to reclaim excreted Mn from bile. Single cell datasets have elucidated that these transporters are not all acting in the same cell types, underscoring the complexity of Mn regulation (5). More than 95% of Mn excreted by the body is via the hepatobiliary route. To date, Mn toxicity has been the focus of study given significant neurologic effects of Mn accumulation (6, 7). Less is known about the physiologic effects of relative Mn insufficiency, although human studies clearly show Mn insufficiency drives disordered glycosylation and mitochondrial dysfunction; in the liver, decreased Mn content has been correlated with fat accumulation in non−alcoholic fatty liver disease, again reinforcing the plausible physiologic disruptions with altered Mn homeostasis (8, 9).

In this work, across multiple models of ‘illness’, we find evidence for Mn fluxes. When dietary Mn is in excess, liver Mn levels decrease while biliary Mn increases. This change in hepatobiliary Mn homeostasis occurs in response to multiple models of acute intestinal inflammation and infection, but also occurred in a model of systemic infection with *Candida albicans*. When dietary Mn intake is only adequate and not in excess, driving down baseline liver Mn, upon induction of colitis, liver Mn did not decrease further, however biliary Mn was still markedly increased. In a model of chronic inflammation in the gut using a pathobiont enterotoxigenic Bacteriodes fragilis, hepatic Mn was decreased and biliary Mn was increased compared to sham treated mice, despite resolution of the acute systemic signs of infection. Finally, we found differential expression of ZIP8 and ZNT10 in in the liver when the mice were tested in the DSS colitis model, but not ZIP14. Taken together, across acute and chronic models of intestinal and extraintestinal ‘illness’, in both male and female mice, and in mice from two different genetic backgrounds, these findings support Mn homeostasis is dynamic. Modeling ‘illness’ in mice provides an important system to uncover mechanisms governing Mn homeostasis and the physiological effects for the host in acute and chronic disease.

## Methods

### Animal studies

Wild−type C57/BL6 mice were used in these studies. Mice were obtained from our colony (WT mice derived from heterozygous breeding of Zip8 393T−KI mice (3)) or purchased from Jackson Labs. We housed the mice in a single facility. We weaned mice onto conventional mouse chow (Harlan/Envigo Teklad Global 18% Protein Extruded Diet 2018SX, 100 ppm Mn) or a purified diet (Envigo, purified TD.97184 to include 10 ppm Mn and ∼40 ppm Fe). Sex and age of mice in the figure legends. On the evening before sacrifice, mice were fasted to facilitate collection of bile. Mice were euthanized using 5% isoflurane. We collected blood via cardiac puncture and reserved it for whole−blood Mn. Gallbladder bile was collected by puncturing the distended gallbladder using a 27G insulin syringe. We harvested organs as previously reported (3).

### Dextran sodium sulfate (DSS) colitis

Male and female mice were exposed to 3% (w/v) DSS (MP Biomedicals; MW = 36,000−50,000 Da) dissolved in sterile water for 3−5 days as previously reported (3). We assessed survival, percent body weight loss and rectal bleeding daily. We sacrificed mice on day 8 or day 14 of the experiment after an overnight fast. We collected whole blood and bile as outlined above. We measured body weight, colon length, cecal weight, and spleen weight to assess the severity of colitis and studied the distal colon for histology (10).

### *Candida albicans* infection

To prepare for infection, clinical isolate SC5314 was grown in YPD (1% yeast extract, 2% peptone, 2% dextrose) at 30°C, 220rpm to stationary phase to an OD_600_ of 15. 11−week old Balb/c mice were injected with 5×10^5^ live *C. albicans* cells by lateral tail vein. Mice were euthanized 72 hours post−infection by cervical dislocation, and liver sections ranging from 100mg−400mg were harvested.

### Enterotoxigenic Bacteroides fragilis (ETBF) colitis

Mice were inoculated with BFT−2−secreting ETBF as previously reported (11). To facilitate infection, we pre−treated animals with clindamycin (0.1g l^−1^) and streptomycin (5 g l ^−1^) for 5 days before peroral ETBF inoculation (∼1×10^8^ bacteria in phosphate buffered saline (PBS)) or PBS alone (sham control). For the chronic model, we confirmed fecal colonization as colony−forming units per g stool. We recorded weight daily for the first week and then weekly to 4 weeks. At sacrifice, we treated the animals as outlined above.

### Atomic absorption spectroscopy (AAS)

We carried out metal analysis of whole blood and bile as previously reported.(3) For the *Candida albicans* infection experiment, liver sections were washed with 1mL of TE buffer (10 mM Tris, 1 mM EDTA, pH 8.0), and twice with 1mL of milli−Q water before digestion. Liver sections were digested in 500μL of 20% nitric acid (Fisher Chemical) at 90°C overnight. Samples were diluted to 2% nitric acid with milli−Q water before Mn concentration was analyzed using a graphite furnace atomic absorption spectrophotometer, AAnalyst 600 (Perkin Elmer).

### Inductively coupled plasma mass spectrometry (ICP−MS)

We carried out metal analysis of liver tissue as previously reported (3).

### mRNA extraction and quantitative PCR

RNA was extracted using the QIAGEN RNeasy Mini Kit per the manufacturer’s protocols. Purified RNA concentrations were measured using wavelengths of 260/280 nm on the Beckman Coulter DU 800 Spectrophotometer. From equal amounts of RNA, cDNA was generated using iSCRIPT. Real−time quantitative PCR was performed and analyzed using Taqman reagents on the QuantStudio 6 flex real time PCR system using specific TaqMan Gene Expression Assays (primers, Mm01320977_m1 Slc39a8, Mm01315481_m1 Slc30a10 Mm01317439_m1 Slc39a14, Mm99999915_g1 Gapdh) ((Thermo Fisher Scientific). Gapdh was included as an internal control, and relative expression was calculated using the 2(–delta delta Ct) method.

### Western blot

Samples (20 mg) of flash−frozen liver tissue were lysed and homogenized with sucrose lysis buffer (50 mM HEPES, 150nM NaCl, 0.25M Sucrose, 0.4nM AEBSF, 1:100 Igepal) with complete protease inhibitor (Roche Diagnostics, Mannheim, Germany). Lysate samples were centrifuged at 15,000 g for 10 min. Protein concentrations were determined using a Bio− Rad protein assay dye reagent (#5000006), Bradford method−based colorimetric assay. Proteins were mixed with Laemmli buffer, incubated at 42 °C for 15 min, electrophoretically separated on a 10% tris glycine Invitrogen gel, and transferred to a nitrocellulose membrane. The blot was blocked for 1 h in 5% (w/v) nonfat dry milk in Tris−buffered saline containing 0.1% (v/v) Tween 20 (TBST). The blots were then incubated overnight at 4 °C in blocking buffer containing rabbit anti−ZIP8 (Proteintech # 20459−1−AP, 1:1000). Mouse anti−GAPDH (Sigma #G8795) was used as loading control. The membranes were washed three times for 10 min with TBST and incubated with the appropriate secondary antibodies. Immunoreactivity was visualized by using Odyssey CLx Imager.

### Statistics

We performed statistical analysis using Prism Graphpad. We plotted mean and standard error of the mean (S.E.M.) with each individual data point shown. When two groups were analyzed, we used Mann−Whitney t−test to accommodate differences in standard deviation between groups. When three groups were analyzed, we used Brown−Forsythe and Welch’s ANOVA with Bennett multiple comparison’s testing again to account for differences in standard deviation. We performed two−way ANOVA to study sex−specific effects in total metal measurements (Fig. 1A). Outlier testing was not performed, therefore no data were removed. We considered a p−value of <0.05 statistically significant, and we indicate statistical significance by the p−value or asterisks as convention.

**Figure 1.**
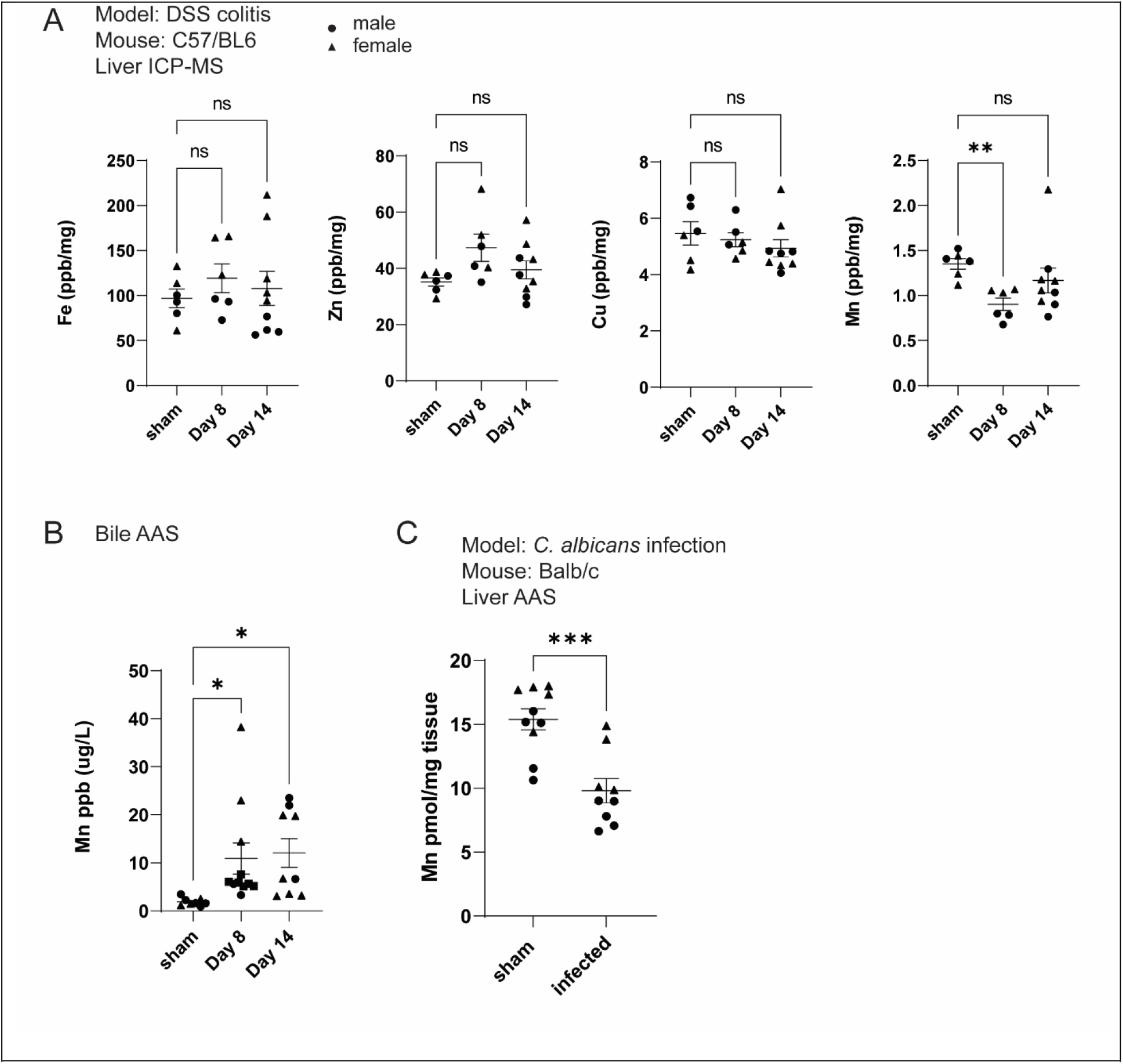
Total liver Mn decreases and biliary Mn increases in the setting of intestinal inflammation. (A) Total liver iron, zinc, copper, and Mn as measured by ICP−MS normalized to dry weight of tissue. C57/BL6 mice were exposed to dextran sodium sulfate (DSS) in the drinking water for 5 days and then allowed to recover with mice sacrificed at day 8 and day 14. (B) Bile Mn as measured by AAS in sham mice and mice exposed to DSS at day 8 and day 14. (C) Total liver Mn measured by AAS. Balb/c mice were infected with *C. albicans* by tail vein injection and sacrificed at 72 hours after infection. N=5−9 mice/group. Sex indicated by circles (males), triangles (females). Statistical significance determined by Brown−Forsythe and Welch’s ANOVA with Bennett multiple comparison’s testing (A, B) and unpaired t−test (C) with significance noted by p−value indicated by one asterisk (<0.05), two asterisks (<0.005) or three asterisks (<0.0005).

### Study approval

Animal experiments were approved by the Johns Hopkins University Animal Care and Use Committee in accordance with the Association for Assessment and Accreditation of Laboratory Animal Care International.

## Results

### ‘Illness’ increases biliary Mn and decreases hepatic Mn in mice fed standard house chow

Acute intestinal inflammation was modeled using the dextran sodium sulfate (DSS) colitis model. Mice were exposed to dextran sodium sulfate (DSS) in drinking water for 5 days. We examined total liver iron, zinc, copper and Mn in the liver by ICP−MS. Mn was the only significantly different metal at the time points studied with total liver Mn decreased at day 8 and returning to near baseline at day 14 (**Fig. 1A**). We tested for sex−specific effects by two−way ANOVA and found only an effect of sex in total iron (p=0.003), but no sex−specific effects were statistically significant with Mn, copper, or zinc.

Because bile is the major site of Mn excretion, we measured biliary Mn. Gallbladder bile was collected from fasted mice at time of sacrifice, and Mn was measured by AAS. Biliary Mn was increased at day 8 and day 14 (**Fig. 1B**). Biliary Mn increased approximately 3−fold (p=0.0166).

Next we asked if this change in hepatic and biliary Mn was in response to intestinal inflammation or if extra−intestinal inflammation or infection would drive the same phenomenon. To address this question, we used an infection model of *Candida albicans*. These experiments also allowed us to study mice of a different genetic background, Balb/c, fed standard house chow. In this model, the mice were infected with *Candida albicans* via tail vein injection and sacrificed at 72 hours after infection. The kidney is the principal site of organ involvement. Consistent with the DSS experiments, hepatic Mn decreased in male and female mice (**Fig. 1C**).

### Increased biliary Mn in response to colitis is maintained when dietary Mn is adequate but not in excess

Mn status and tissue levels of Mn are determined by dietary Mn intake. We next studied the impact of decreased dietary Mn on hepatobiliary Mn excretion in mice treated in the DSS colitis model. The mice in our facility are fed conventional, corn−based chow that contains at least 10 times more Mn than considered adequate for mice (Mn 100 ppm) (12). We sought to study the effect of a diet with a moderated, yet adequate Mn content (Mn 10 ppm) permitted by a purified dietary background. Male and female C57/BL6 mice were weaned onto the purified diet and fed for 8−12 weeks prior to experiments. The purified diet decreased baseline hepatic Mn levels by approximately 60% as expected with decreased dietary Mn (**Fig. 2A**, p=0.0095, Mann−Whitney). We again challenged mice in the DSS colitis model. Demonstration of induction of colitis is shown by body weight ratio over time course, colon, cecum, and spleen size at sacrifice, and representative histology (**Fig. 2B−D**). There was significant lethality leaving only sufficient mice to analyze at the day 8 time point. Here, starting at the lower baseline Mn, total liver Mn was not statistically different at day 8 (**Fig. 2E**), however, the increase in biliary Mn was almost 20−fold higher (**Fig. 2F**, p=0.0159, Mann−Whitney). These data suggest that even in the setting of decreased dietary Mn intake using a purified diet that decreases baseline liver Mn, hepatobiliary Mn excretion remains increased in male and female mice in the acute DSS colitis model.

**Figure 2.**
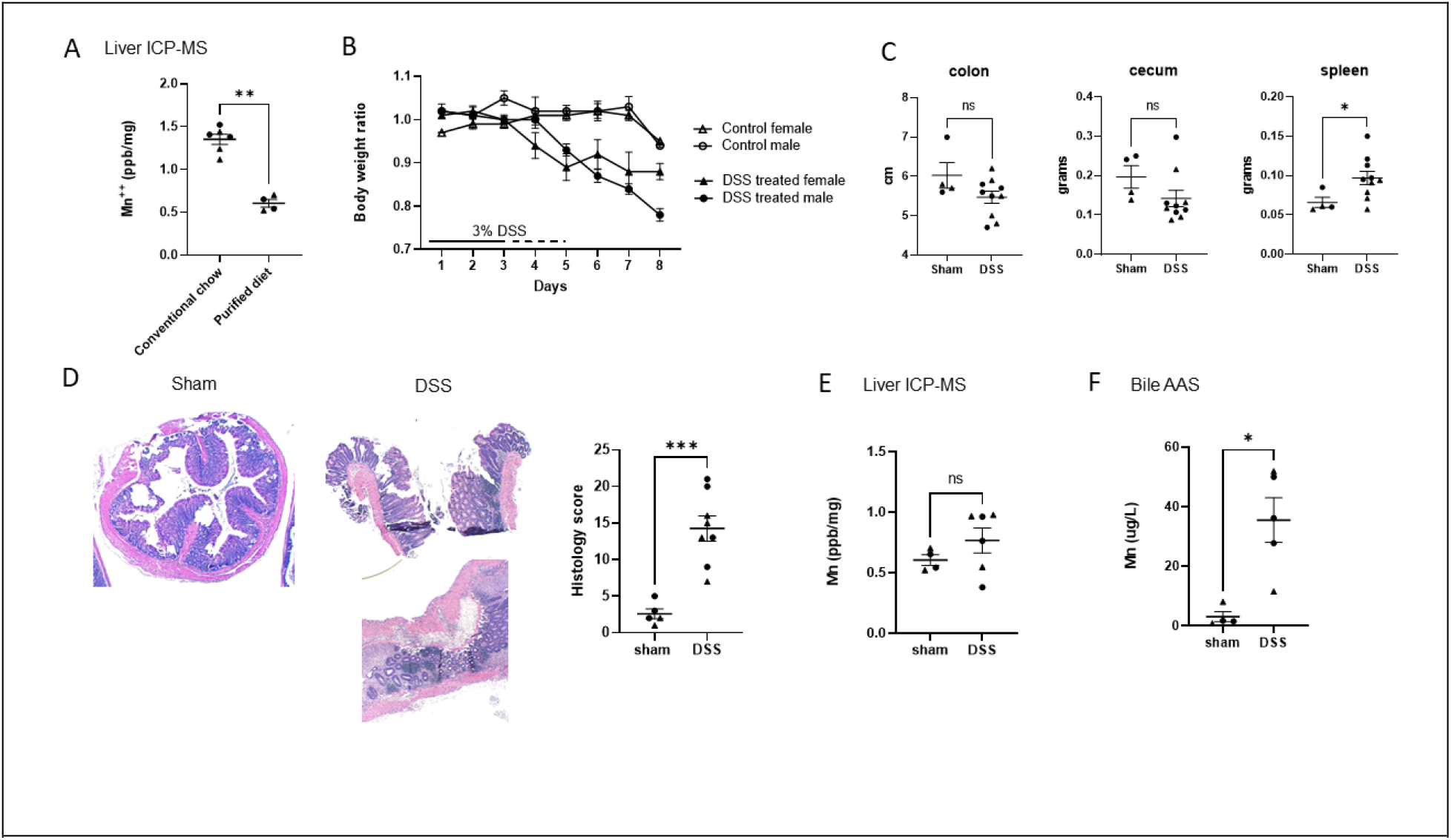
Increased biliary Mn in response to colitis is maintained on purified diet containing Mn 10 ppm. (A) Baseline total liver Mn of WT mice fed conventional chow and purified diet. (B) Body weight ratio over DSS colitis time course in sham (control) and DSS treated mice. The experiment was repeated twice with 3% DSS in drinking water for 3−5 days. (C) Colon length, cecal size, and spleen size at time of sacrifice. (D) Representative histology from sham and DSS−exposed mice showing evidence of severe colitis with ulceration, elongation of crypts, and transmural inflammation and histology score (10). (E) Total liver Mn of sham versus DSS exposed mice fed purified diet containing Mn 10 ppm (ICP−MS). (F) Bile Mn of sham and DSS exposed mice (AAS). Body weight curves incorporate data across multiple experiments with N=4 (sham) and N=6−7 mice (treated) in each experiment. Representative data in figures 2C, 2D−F with individual data points graphed. Mean and S.E.M. is graphed with statistical analysis performed by Mann−Whitney with p−value indicated by one asterisk (<0.05), two asterisks (<0.005), or three asterisks (<0.0005).

### Despite recovery from acute infection, increased biliary Mn persists in a second model of colitis with chronic inflammation

To replicate our findings in a second model of intestinal inflammation, we used an infection model with the Crohn’s disease−associated pathobiont, *enterotoxigenic Bacteroides fragilis* (ETBF) (11). In addition to a high prevalence in patients with Crohn’s disease, ETBF has been reported to colonize healthy adults (up to 40% in one small study) and may contribute to long− term colon cancer risk (11, 13). In the animal model of ETBF infection, WT mice were fed the conventional house chow, pre−treated with antibiotics and then infected with ETBF and followed for 27 days as previously reported (**Fig. 3A**) (11). Consistent with an acute colitis, mice lose weight to approximately 7 days and then enter a phase of chronic infection (**Fig. 3B**). Despite weight recovery, at sacrifice at day 27, the infected mice have contracted cecums and enlarged spleens (**Fig. 3C**). These findings are consistent with an ongoing, mild inflammatory response as previously reported (11). At day 27, total liver Mn remains decreased (**Fig. 3D**), and biliary Mn remains increased (**Fig. 3E**). Blood Mn was also decreased in ETBF−exposed mice (**Fig. 3F**). These data confirm that hepatobiliary Mn homeostasis remains perturbed in a model of chronic colonic infection.

**Figure 3.**
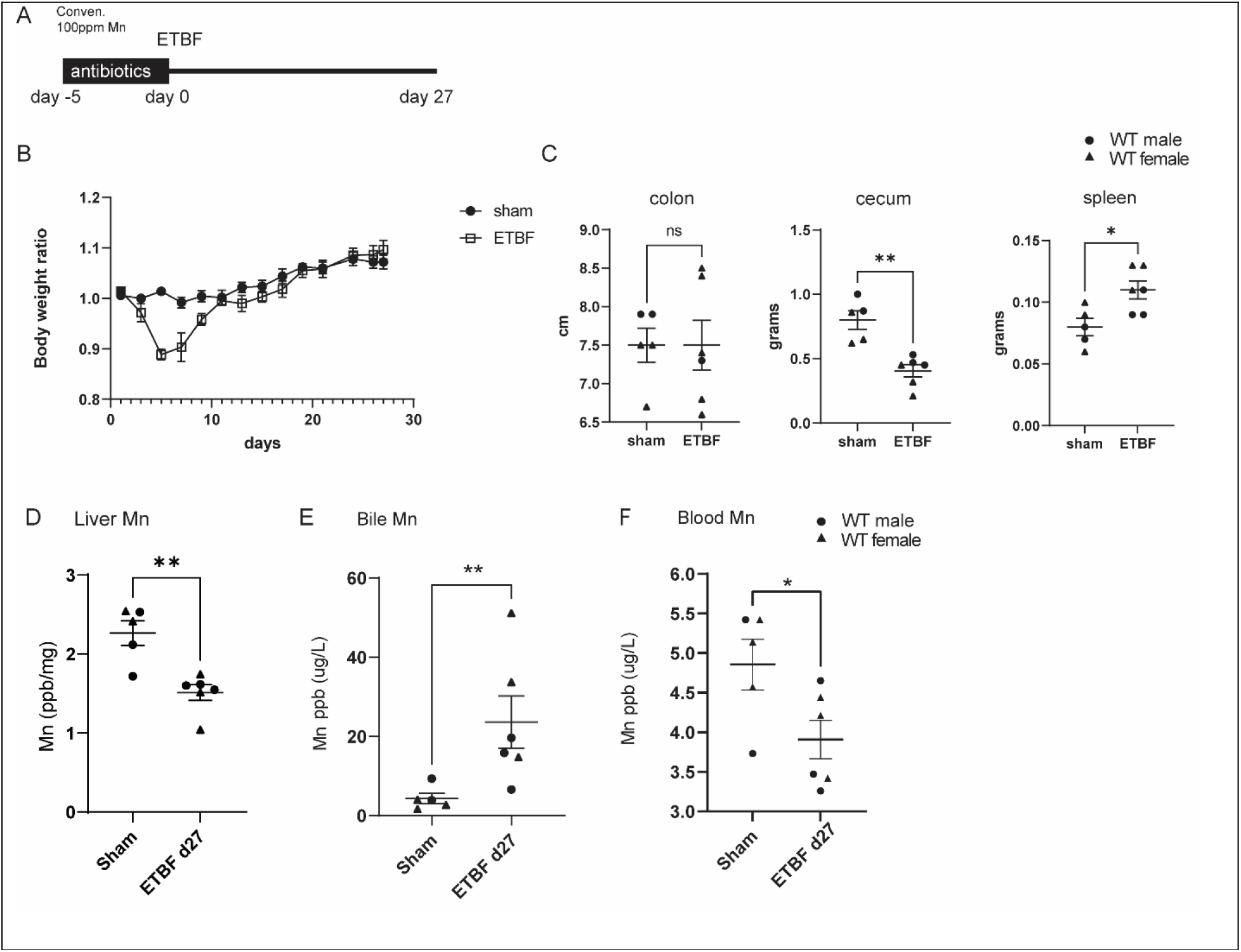
Biliary Mn is increased in a second model of intestinal inflammation, *enterotoxigenic Bacteroides fragilis* (ETBF) acute and chronic infection. (A) Schematic of model depicting pre−treatment with antibiotics at day −7, ETBF oral inoculation at day 0, and sacrifice at day 27. (B) Body weight ratio over time course of experiment determined by mouse weight at time point divided to its own day 1 weight. (n=5−6 mice/group) (C) At sacrifice at day 27, colon length, cecum weight, and spleen weight were measured. (D) Liver Mn is decreased at day 27, AAS. (E) Biliary Mn is increased at day 27 in ETBF−infected mice as measured by AAS. (F) Blood Mn is decreased at day 27 in ETBF−infected mic. Individual data points (with sex indicated in legend), mean and S.E.M. graphed with statistical significance determined by Mann−Whitney with p−value indicated by one asterisk (<0.05) or two asterisks (<0.005). N=5−6 mice/group.

### Mn transporters *Slc39a8*/Zip8 and *Slc30a10*/Znt10 are differentially expressed in the liver in response to intestinal inflammation

Finally, given the evidence for dynamic Mn homeostasis, in DSS−treated wild−type C57/BL6 mice, we performed qPCR on total liver mRNA. As we reviewed in the introduction, systemic and hepatic Mn homeostasis is regulated by the activity of three transporters – *Slc49a14*/Zip14 (basolateral uptake of Mn into hepatocytes), *Slc39a8*/Zip8 (apical Mn uptake from bile) and *Slc30a10*/ZnT10 (apical Mn efflux into bile) (4). *Slc39a8* and *Slc30a10* mRNA were both decreased at day 9 (4 days after DSS withdrawal) and day 14 compared to sham animals, while *Slc39a14* was unchanged (**Fig. 4A**). At the protein level, Zip8 expression was decreased at day 9 with trend toward recovery at day 14 (**Fig. 4B)**. Decreased Zip8 expression will decrease the amount of Mn reclaimed from bile and result in increased biliary Mn content as observed in these experiments, especially at the early time points (14). Integrating the mRNA−level data for *Slc30a10* appears to not support an increase in active Mn export to account for the marked increase in biliary Mn, but this will require further study.

**Figure 4.**
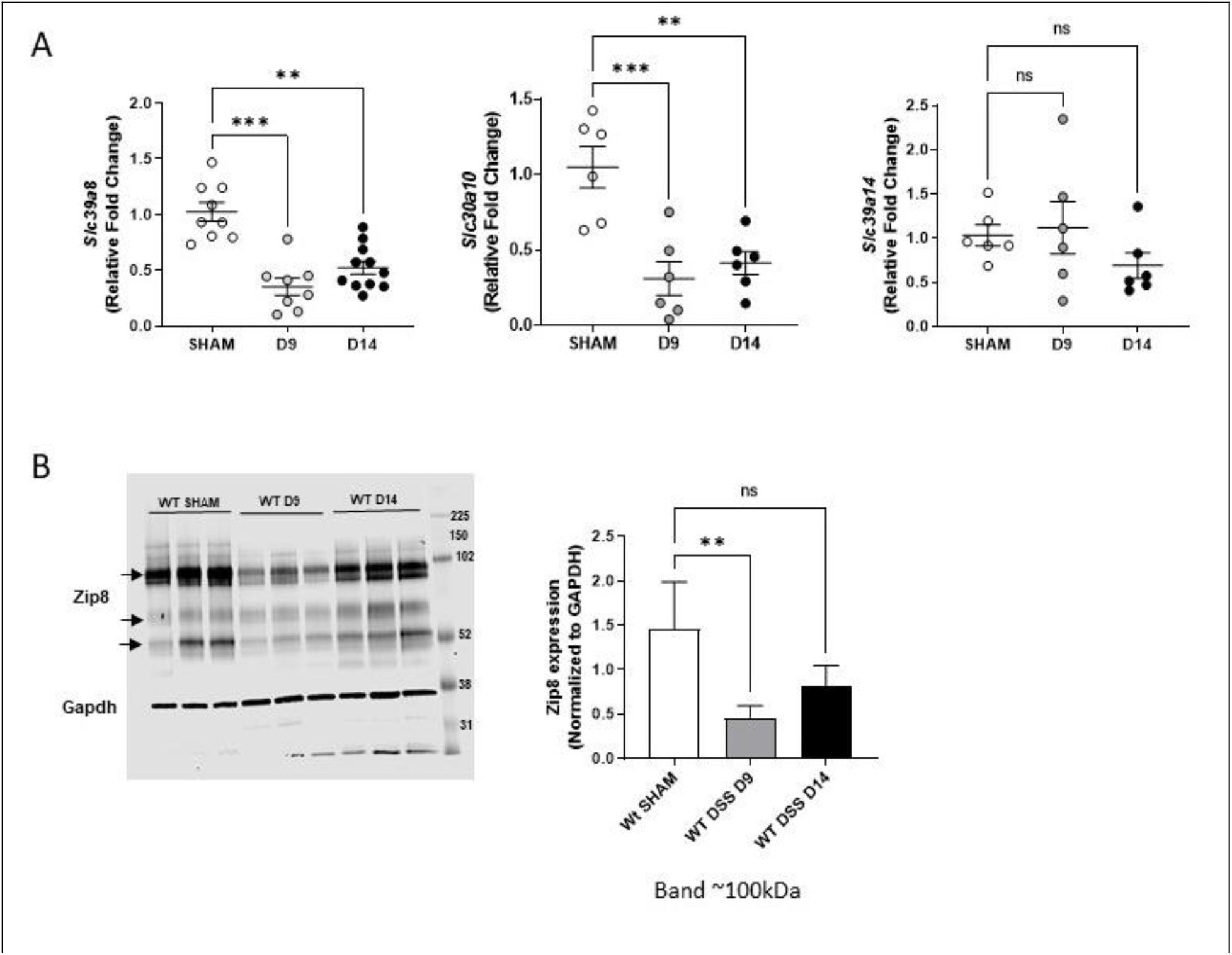
Zip8 protein decreases in response to intestinal inflammation. (A) qPCR of mRNA isolated from C57/BL6 WT mouse liver at day 9 or day 14 of DSS colitis model to study *Slc39a8, Slc30a10*, and *Slc39a14* expression. (B) Western blot for Zip8 in sham and WT mice in the DSS colitis model with decreased Zip8 expression by immunoblot and densitometry. Zip8 is represented by multiple bands on immunoblot: monomeric bands at ∼52kDa (unglycosylated Zip8), ∼76 kDa (glycosylated Zip8) and ∼100kDa (oligomeric, glycosylated Zip8 bands) from total liver protein lysate. N=5−6 mice, C57/BL6 male mice.

## Discussion

This study demonstrates that Mn homeostasis is dynamic in response to illness in mice. Illness– here studied using multiple acute and chronic models − induces a relative hepatic insufficiency (when dietary Mn is in excess) and increased biliary Mn regardless of baseline Mn status. This study prioritizes further elucidation of the physiological effects of relative decreases in liver Mn and increases in biliary Mn.

Our data build on a small body of literature in human and mice studies that have suggested Mn homeostasis is dynamic and responsive to systemic drivers, including hormones. In one study, mice exposed to high doses of prednisolone or cortisol retained less radiolabeled Mn in liver than controls (15). A second study examined turnover of Mn in patients with rheumatoid arthritis. Uptake of injected ^54^Mn did not differ between arthritis patients and disease control patients (with diagnoses including diabetes, Friedreich’s ataxia, cerebral vascular thrombosis), but the biological half−life was longer in arthritis patients, consistent with slowed turnover (16). The findings of this report are confounded by an interaction between Mn clearance and medications, but both studies support this overarching concept that Mn metabolism is responsive to systemic disease.

The detrimental effects of Mn toxicity have long been appreciated, especially in the context of environmental exposures (6). However, there is accumulating evidence that greater focus on the effects of Mn insufficiency is necessary. This is particularly relevant to our findings that hepatic Mn is decreased in setting of ‘illness’ when the mice are fed a chow with Mn in excess. It is thought that humans typically consume more Mn than required, and Mn deficiency is rare, but there is evidence that dietary Mn intake has declined by approximately 75% in the last century (1, 17). Recent work has shown that hepatic Mn is decreased in earlier stages of chronic liver disease, and there is a strong inverse relationship between levels of hepatic Mn and severity of steatosis with lower levels of Mn correlating with higher steatosis scores (8). Further, a pathogenic mutation in the Mn transporter ZIP8 (A391T; rs13107325) that decreases liver Mn and increases biliary Mn has been identified as one of the most pleiotropic variants in the human genome (3, 18, 19). ZIP8 391−Thr is associated with obesity and early−childhood− onset obesity, schizophrenia, and Crohn’s disease, while protective against alcohol misuse disorder and hypertension (20). ZIP8 391−Thr and a second variant in the Mn exporter, ZNT10, were two of six genetic variants associated with an MRI−based, non−invasive measure of steatohepatitis and fibrosis, and the variants in ZIP8 and ZNT10 were both associated with elevated aminotransferases in the UK Biobank (21). These epidemiological and genetic studies emphasize the potential important clinical implications of aberrant Mn homeostasis in the liver.

If there are physiologic effects of increased biliary Mn is not understood. Fundamentally, is biliary excretion of Mn simply a means of disposal to regulate systemic Mn availability or does it have its own physiologic significance? It is very likely to depend on the bioavailability of biliary Mn both to the host and the resident microbes, but this has not been explored. Recently, ileal Mn excretion via Znt10 was shown to be dependent on bile acid composition, suggesting that there is an interplay between Mn homeostasis and bile acids, but how this connects to biliary Mn fluxes in illness remains to be clarified (22). In our data, unexpectedly, biliary Mn was the most increased when the mice were fed the Mn−adequate purified chow that decreased baseline hepatic Mn levels. These animals were the most sick of any of the ‘illness’ models, and, in addition to Mn levels, the fiber source differs between purified diets and conventional, corn− based chow. Therefore, it will be necessary in future studies to dissect if biliary Mn excretion relates to disease severity or other dietary factors, in addition to Mn.

Our transcriptional and protein data support down−regulation of Zip8 as a lead cause of increased biliary Mn in response to ‘illness.’ Regulation of ZIP8 expression in response to inflammation is well−established in the literature with examples of both increased and decreased expression: ZIP8 expression (mRNA and protein) increases in macrophages (mouse and human) and the lung (mouse) in response to LPS, primarily through NF−κB mediated transcriptional regulation and affecting zinc uptake (23). However, consistent with our data, *Slc39a8* mRNA expression has been reported to be decreased in the liver upon systemic LPS exposure in a mouse model (24).

The major strength of this study is the utilization of multiple models of intestinal inflammation and infection, a model of extra−intestinal infection (*Candida albicans*), testing in male and female mice, and testing in wild−type mice of two genetic backgrounds to confirm the finding of dynamic Mn homeostasis in response to illness. This present study remains descriptive, and future studies need to focus on how the liver ‘senses’ Mn status and what ‘alarm’ signals might induce change in Mn homeostasis during illness. It will be important to establish if differential Mn flux affects host fitness, examine how the relative hepatic Mn insufficiency impacts Mn−dependent processes, including glycosylation, redox homeostasis, and gluconeogenesis, and if there is any impact of change in biliary volume or flow rate in the setting of intestinal inflammation. Future studies should dissect the interaction between ZIP8 and other known Mn transporters that participate to hepatic Mn homeostasis, including ZIP14 and ZNT10, in the setting of inflammation (24).

## Acknowledgments

The authors acknowledge the helpful advice of Drs. Shaoguang Wu, Xinqun Wu, and Cynthia Sears. We acknowledge the technical assistance and core resources of the NIH/National Institute of Diabetes and Digestive and Kidney Diseases Johns Hopkins Conte Gastrointestinal Core Center (P30 DK−089502), Department of Pharmaceutical Sciences, University of Maryland (ICP−MS; Michel Lab, NSF CHE1708732), Lutsenko Lab, Johns Hopkins University, Department of Physiology (AAS), and Dr. Brendan Cormack, Johns Hopkins University Molecular Biology and Genetics (*C. albicans* infection model). Funding was provided by the NIH (DK114478 to JM; R21 AI154726 to VC), pilot funding from Johns Hopkins Specialized Center for Research Excellence in Sex Differences (U54AG062333), and American Gastroenterological Association (JM).

## Author contributions

**Laxmi Sunuwar:** conceptualization, methodology, validation, analysis, visualization, writing, reviewing, editing. **Vartika Tomar:** validation, analysis, visualization, writing, reviewing, editing. **Asia Wildeman:** methodology, validation, analysis, visualization, writing, reviewing, editing. **Valerie Culotta:** methodology, validation, analysis, visualization, writing, reviewing, editing. **Joanna Melia:** conceptualization, methodology, analysis, resources, writing the first draft of the manuscript, reviewing, editing, visualization, supervision, and funding acquisition.

